# SPARC promotes cell invasion in vivo by decreasing type IV collagen levels in the basement membrane

**DOI:** 10.1101/037218

**Authors:** Meghan A. Morrissey, Ranjay Jayadev, Ginger R. Miley, Catherine A. Blebea, Qiuyi Chi, Shinji Ihara, David R. Sherwood

## Abstract

Overexpression of SPARC, a collagen-binding glycoprotein, is strongly associated with tumor invasion through extracellular matrix in many aggressive cancers. SPARC regulates numerous cellular processes including integrin-mediated cell adhesion, cell signaling pathways, and extracellular matrix assembly; however, the mechanism by which SPARC promotes cell invasion *in vivo* remains unclear. A main obstacle in understanding SPARC function has been the difficulty of visualizing and experimentally examining the dynamic interactions between invasive cells, extracellular matrix and SPARC in native tissue environments. Using the model of anchor cell invasion through the basement membrane (BM) extracellular matrix in *Caenorhabditis elegans,* we find that SPARC overexpression is highly pro-invasive and rescues BM transmigration in mutants with defects in diverse aspects of invasion, including cell polarity, invadopodia formation, and matrix metalloproteinase expression. By examining BM assembly, we find that overexpression of SPARC specifically decreases levels of BM type IV collagen, a crucial structural BM component. Reduction of type IV collagen mimicked SPARC overexpression and was sufficient to promote invasion. Tissue-specific overexpression and photobleaching experiments revealed that SPARC acts extracellularly to inhibit collagen incorporation into BM. By reducing endogenous SPARC, we also found that SPARC functions normally to traffic collagen from its site of synthesis to tissues that do not express collagen. We propose that a surplus of SPARC disrupts extracellular collagen trafficking and reduces BM collagen incorporation, thus weakening the BM barrier and dramatically enhancing its ability to be breached by invasive cells.

## Author Summary

SPARC is an extracellular matrix protein that is present at high levels in many metastatic cancers where it promotes tumor invasion into neighboring tissues. The mechanism linking a surplus of SPARC to cell invasion, however, is not clear due to the challenge of examining SPARCs function in complex tumor environments. We have used anchor cell invasion in *C. elegans* development to understand how an excess of SPARC promotes invasion in a native tissue setting. Anchor cell invasion allows experimental examination and visualization of the interactions between an invasive cell, the surrounding tissues, and the basement membrane, a sheet-like extracellular matrix that surrounds tissues. We find that increased SPARC expression potently enhances the ability of weakly invasive anchor cells to breach the basement membrane. Our data indicate that SPARC functions normally to transport the basement membrane component type IV collagen between tissues to precisely regulate its deposition into basement membranes. Collagen molecules are covalently cross-linked and provide basement membranes their barrier properties. Our results indicate that overexpression of SPARC interferes with collagen trafficking and significantly decreases collagen incorporation into basement membranes, thus weakening this barrier and allowing it to be more easily breached by invasive cells.

## Introduction

Cell invasion through extracellular matrix (ECM) is a critical step in the progression of many diseases including cancer metastasis (1, 2). Basement membrane (BM), a dense, sheet-like form of ECM that surrounds most tissues, poses a significant barrier for invasive cells (3). Although traditionally thought of as a static scaffold, changes to ECM composition, crosslinking, and stiffness can promote tumor progression by altering cell survival, growth, migration, or invasion (4, 5). How alterations in BM composition influence cell invasion is not well understood due to the experimental inaccessibility of cell invasion events *in vivo* and the difficulty of replicating native BM and the surrounding microenvironment *in vitro* (1).

The BM is constructed on a cell-associated polymeric laminin scaffolding overlaid with a type IV collagen network (6). BMs also contain a number of associated proteins that link these networks, including nidogen and perlecan (6). Helical collagen trimers assemble into a network through end-on and lateral attachments, including intermolecular covalent cross-links (1, 7, 8). Eliminating collagen α1 and α2 in mice results in the formation of unstable BMs that disintegrate during increased mechanical demands on tissues prior to birth, highlighting the crucial role of collagen in maintaining structural integrity of the BM (9). Collagen is also required for BMs to provide a constrictive force that shapes tissue architecture (10-13). How the large collagen trimers (approximately 400 nm (14)) are trafficked and targeted to the BM is not well understood.

Secreted Protein Acidic and Rich in Cysteine (SPARC) is a collagen-binding matricellular glycoprotein that associates with BM (15, 16). SPARC is overexpressed in many types of human cancers and is a marker of poor prognosis (17). For example, SPARC overexpression has been correlated with tumor metastasis in breast invasive ductal carcinoma, glioblastoma, pancreatic ductal adenocarcinoma, clear-cell renal cell carcinoma, melanoma, and prostate carcinoma (reviewed in (18, 19)). Supporting a direct role for SPARC in promoting metastasis, murine breast and pancreatic cancer metastasis models have shown that elevated SPARC levels increase metastatic potential (20, 21). Furthermore, increased SPARC or exogenous addition of SPARC enhances invasion in *in vitro* assays (20, 22-27). Although a crucial factor in promoting cancer metastasis and invasion, the mechanism(s) by which increased SPARC expression enhances invasive ability *in vivo* is not understood (18, 19).

SPARC has been implicated in numerous cellular processes, including growth factor signaling, cell-ECM interactions, and ECM assembly (18). Several of these putative SPARC functions may promote cell invasive behavior by facilitating BM breach, including a positive correlation between SPARC and matrix metalloproteinase (MMP) expression (28-31) or integrin signaling (32-36). SPARC also plays a conserved role in regulating assembly of BM type IV collagen and interstitial fibrillar collagen (37-41); and may affect ECM by modulating collagen processing (32, 42) or through a direct chaperone-like interaction with type IV collagen (43-45). In addition, SPARC has been proposed to regulate ECM disassembly (46). As the majority of mechanistic studies examining SPARC function during cancer metastasis have been conducted using *in vitro* cell culture models, it is not clear which, if any, of these mechanisms are relevant to SPARCs pro-invasive function *in vivo.*

Anchor cell (AC) invasion during *C. elegans* larval development is an experimentally tractable model to mechanistically examine BM remodeling, BM dynamics, and BM transmigration *in vivo* (3, 47-49). AC invasion through the juxtaposed gonadal and ventral BMs is highly stereotyped and occurs in tight coordination with the divisions of the underlying vulval precursor cell (VPC) P6.p. The AC breaches the BM using F-actin rich invadopodial protrusions after P6.p has divided once (P6.p two-cell stage, Fig 1A), and completely removes the underlying BM after a second round of VPC divisions (P6.p four-cell stage; ˜ 90 minute time period). Following invasion, the VPCs divide again and invaginate (P6.p eight-cell stage). Previous research has identified several pathways that promote distinct aspects of AC invasion (summarized in Fig 1B). The conserved Fos family transcription factor FOS-1A regulates the expression of genes, including the matrix metalloproteinase *zmp-1,* that promote BM breaching (49, 50). Integrin signaling *(ina-1* and *pat-3* heterodimer) stabilizes F-actin regulators at the AC-BM interface that mediate the formation of invadopodia—small, dynamic cellular protrusions that breach the BM (49). Netrin signaling *(unc-6*/netrin and the receptor *unc-40/DCC)* is activated at the site of BM breach and facilitates the generation of a large invasive protrusion that rapidly clears an opening through the BM (51, 52). In addition, AC invasion is stimulated by an unidentified diffusible cue from the underlying vulval cells that activates the Rho GTPase CDC-42, which induces robust invadopodia formation (53-56). Loss of these pathways results in highly penetrant blocks or slowing of AC invasion. Importantly, Fos transcription factors, integrin, netrin, and Cdc42 are all strongly implicated as potent promoters of metastasis and tumor cell invasion (57-62), suggesting that the mechanisms that control AC invasion are conserved and co-opted by tumors (63). Whether SPARC overexpression promotes invasive ability in the AC, similar to its effect on tumor cells in metastatic cancer, is not known.

**Figure 1.**
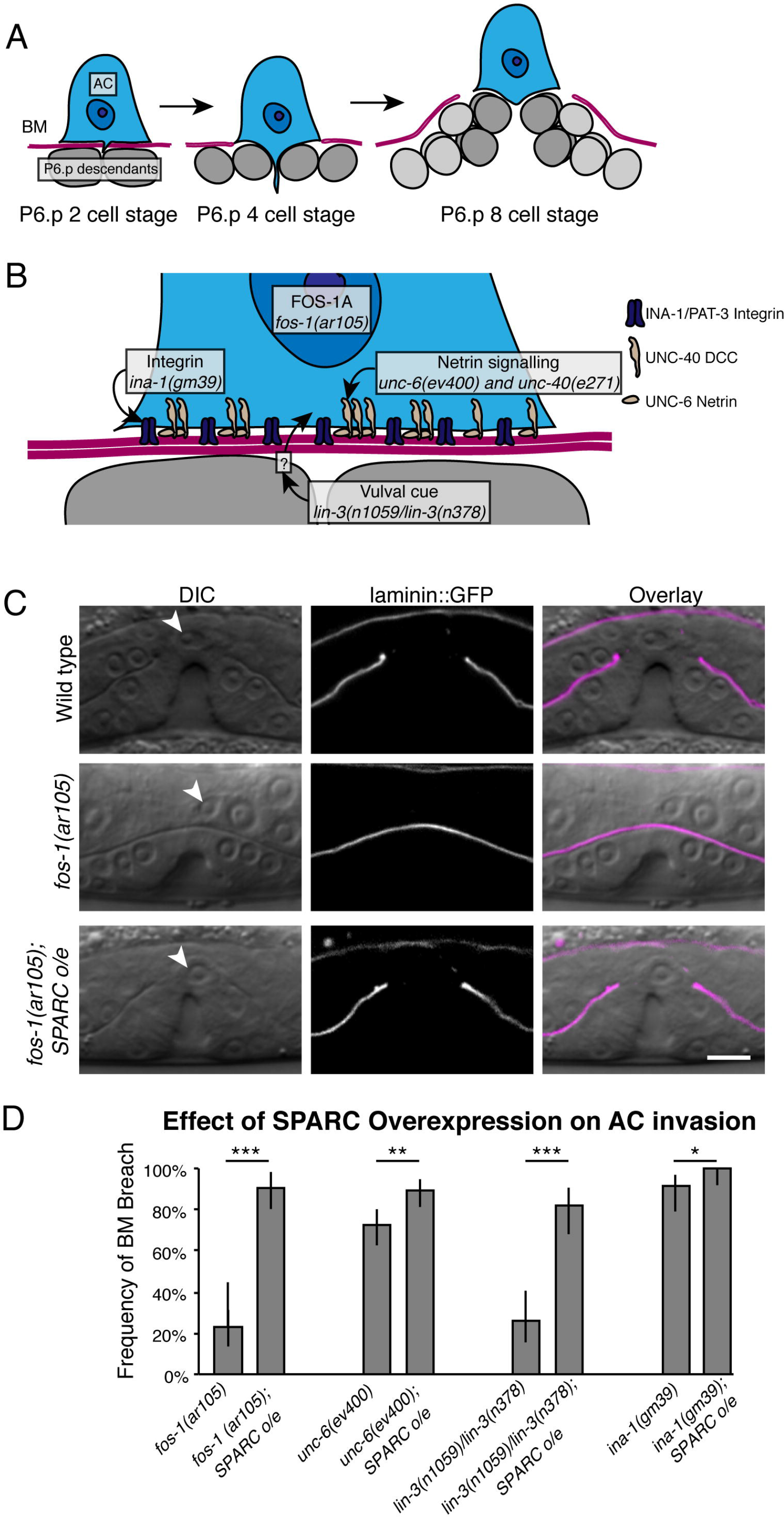
SPARC overexpression promotes AC invasion. (A) A schematic diagram of AC invasion. The AC (blue) breaches the BM (magenta) after the vulval precursor cell P6.p (dark grey) has divided once (P6.p two-cell stage), and completely clears the BM separating the AC and the vulval cells before a second round of P6.p divisions (P6.p four-cell stage). The vulval cells (grey) continue to divide and invaginate (P6.p eight-cell stage). (B) A schematic diagram depicts the major signaling pathways regulating AC invasion. FOS-1A regulates transcription of several pro-invasive genes. Integrin (purple) and netrin (beige) signaling promote F-actin-mediated invasive machinery formation. The vulval cells send an unidentified cue to the AC (absent in *lin-3(n1059)/lin-3(n378)* mutants lacking specification of the vulval cell fate) to stimulate invasion. (C) Representative DIC images overlaid with laminin::GFP fluorescence (magenta) show successful AC (white arrow head) invasion in a wild type animal (top panels); failure of AC invasion in a *fos-1(ar105)* animal (middle); and rescued invasion in a *fos-1(ar105); SPARC overexpression (o/e; syIs113)* animal (lower). (D) The graph depicts the frequency of AC-initiated BM breach at the P6.p eight-cell stage in animals with mutations in different aspects of the AC invasion program with and without SPARC overexpression *(syIs113;* n≥48 for each condition). Error bars show 95% confidence intervals with a continuity correction. ⋆ denotes p<0.05, ⋆⋆ denotes p<0.005, and ⋆⋆⋆ denotes p<0.0005 by two-tailed Fisher’s exact test. Scale bar denotes 5 μm.

In this study we use a combination of genetics, transgenics, and *in vivo* live cell imaging to determine if SPARC overexpression promotes AC invasion. We show that SPARC overexpression restores AC invasion when *fos-1a,* integrin, netrin signaling, or the vulval cue are compromised, indicating SPARC overexpression is strongly pro-invasive. We find that SPARC’s pro-invasive function is dependent on its collagen binding ability and that SPARC overexpression reduces type IV collagen levels in the BM. Decreasing collagen in the BM mimicked the pro-invasive activity of overexpressed SPARC, suggesting that SPARC promotes invasion by reducing BM type IV collagen. Using tissue-specific expression and fluorescence recovery after photobleaching (FRAP), we show that overexpressed SPARC functions extracellularly to reduce the rate of collagen addition into BMs. Further, we find that loss of SPARC during development results in defects in extracellular trafficking of type IV collagen from sites of synthesis to tissues that do not express collagen. Taken together, our results suggest that SPARC directly regulates extracellular type IV collagen transfer between tissues, and that overexpression of SPARC perturbs its normal trafficking function, thus decreasing BM collagen deposition, weakening these barriers and promoting BM transmigration by invasive cells.

## Results

### SPARC overexpression promotes AC invasion in multiple mutant backgrounds

SPARC overexpression is strongly associated with tumor metastasis in many different cancers and promotes cancer cell invasion (18, 19, 64). We thus wanted to determine if SPARC overexpression might also promote AC invasion in *C. elegans.* We created integrated multi-copy transgenes using the endogenous SPARC promoter to drive SPARC (the gene *ost-1* in *C. elegans)* mRNA and protein production at a two-to-five fold increase over normal levels (S1 Fig). As the timing of AC invasion is precisely regulated by multiple extracellular cues and intrinsic factors (47), an increase in SPARC is unlikely to accelerate AC invasion. Consistent with this prediction, ACs in worms overexpressing SPARC completed invasion at the normal time during the P6.p four-cell stage (see Fig 1A; n=27/27 P6.p four-cell stage, 56/56 P6.p eight cell stage animals). Thus, overexpression of SPARC does not appear to alter the endogenous invasion program.

We next asked if SPARC overexpression promotes invasion in mutant backgrounds where invasion is inhibited or slowed by loss of specific pathways. We first examined animals with a null mutation in the transcription factor *fos-1a (fos-1(ar105)).* FOS-1A promotes BM breaching, controlling expression of matrix metalloproteases that degrade ECM (1, 65) and loss of *fos-1a* results in a highly penetrant block or delay in AC invasion (50). While ACs in *fos-1(ar105)* mutant worms successfully breached the BM in only 23% of animals by the P6.p eight-cell stage (n=23/100 animals, Fig 1C and D), overexpressing SPARC dramatically increased the frequency of BM breach to 90% (n=57/63, p<0.0005). Thus, SPARC overexpression promotes invasion in the absence of FOS-1A.

SPARC overexpression may specifically compensate for the absence of FOS-1A or might act independently of this pathway. To distinguish between these possibilities, we overexpressed SPARC in animals harboring mutations in other pathways regulating AC invasion - integrin, netrin signaling, and the vulval cue (the latter absent when the vulval cells are not specified in *lin-3(n1059)/lin-3(n378)* worms; Fig 1B (51-53)). Strikingly, in all genetic backgrounds, SPARC overexpression significantly increased the frequency of AC invasion (Fig 1D). These studies indicate that SPARC overexpression is broadly pro-invasive and can compensate for the loss of multiple distinct pathways that promote AC invasion.

### The pro-invasive function of SPARC is dependent on collagen binding

We next sought to elucidate the mechanism by which SPARC enhances invasion. Previous work has indicated that SPARC can alter a number of genes and pathways that could promote invasion *in vivo,* including MMP expression and activation (28-31), as well as integrin signaling (32-36). SPARC overexpression promoted invasion in animals with defects in invadopodia assembly (vulval cue/integrin/*cdc-42),* invasive protrusion formation (netrin pathway), and MMP expression *(fos-1a*), suggesting that SPARC has a broad and possibly distinct function in augmenting the ability of the AC to invade. Previous studies have also implicated SPARC as an important regulator of type IV collagen deposition in BM (12, 39, 41). As type IV collagen is a major structural barrier that cells must overcome to breach BM (1), we hypothesized that SPARC-mediated alterations of BM collagen might allow even weakly invasive ACs to transmigrate the BM barrier. We thus examined the relationship between SPARC and type IV collagen during AC invasion. In *C. elegans* overexpressed SPARC::GFP and endogenous SPARC are produced primarily in the body wall and vulval muscles, and localize to the BMs surrounding all other tissues (Fig 2A and B, (66)). Type IV collagen also localizes to the BM and is expressed primarily in the body wall muscles, with the neuronal ring ganglion, spermatheca, uterine muscles, vulval muscles, and the distal tip cell of the somatic gonad also contributing collagen during specific larval stages (Fig 2A and B; (67)). Previous *in vitro* studies have identified a collagen-binding pocket in SPARC that is highly conserved from *C. elegans* to humans (37, 43, 45, 68, 69). To determine if a direct interaction between SPARC and collagen is required for SPARC to promote invasion, we created two point mutations, R152L and Q159A, in the collagen-binding pocket of SPARC. These point mutations abrogate collagen binding without altering SPARC conformation (43). In addition, these mutations do not disrupt the follistatin domain (residues 52-75), which is sufficient for SPARC to directly bind and activate integrin (70). We drove wild type SPARC and the form of SPARC unable to bind collagen (SPARC^R152L,Q159A^) under the control of a heat shock promoter in the *fos-1(ar105)* mutant background. We found that overexpressed SPARC^R152L,Q159A^ was localized to the BM at levels sufficient to promote invasion (S2 Fig); however, this mutant form of SPARC was not pro-invasive (Fig 2C). In contrast, wild type SPARC driven by the heat shock promoter almost fully rescued AC invasion in the *fos-1* mutant background (Fig 2C). These results indicate that a direct interaction between SPARC and collagen is essential for SPARC’s ability to promote AC invasion.

**Figure 2.**
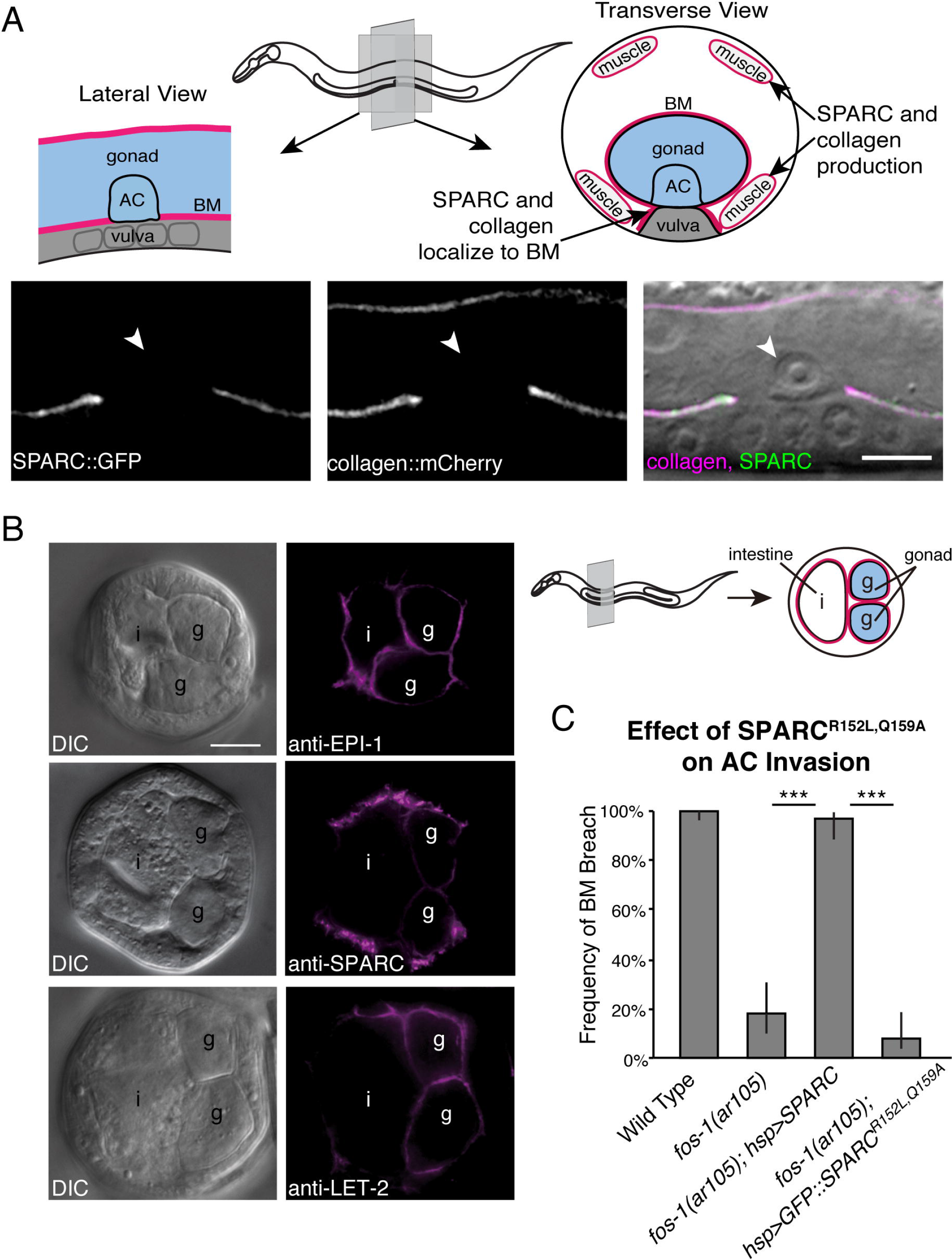
SPARC and type IV collagen colocalize in the gonadal BM and the collagen-binding pocket of SPARC is required to promote invasion. (A) In the top panel, a schematic diagram shows the location of the BM, AC, and vulval cells in a lateral section of the worm (left) and a transverse section (right). The transverse section also includes the body wall muscles, which is the predominant site of type IV collagen and SPARC expression. Below, a lateral section shows SPARC::GFP *(syIs115;* left) and collagen::mCherry (center) fluorescence at the BM overlaid with a DIC image of an invaded AC (arrowhead; right). (B) Immunocytochemistry on fixed transverse worm sections shows that endogenous SPARC (anti-SPARC) and type IV collagen (anti-LET-2) localize with laminin (anti-EPI-1) in the BM. The schematic on the right indicates the approximate position of these sections and the location of the gonad (g) and intestine (i). All proteins colocalize in the gonadal BM. Type IV collagen and laminin localize uniformly in the gonadal BM, whereas SPARC appears to be more strongly enriched in the BM on the outer surface of the gonad. (C) The graph depicts the frequency of BM breach in animals overexpressing SPARC (*hsp>SPARC*) compared to animals overexpressing a form of SPARC *(hsp>SPARC^R152LQ159A^::GFP)* that is unable to bind type IV collagen (n≥51 for each genotype). Error bars denote 95% confidence intervals with a continuity correction. ⋆⋆⋆ denotes p<0.0005 by two-tailed Fisher’s exact test

### SPARC overexpression decreases levels of type IV collagen in the BM

We next wanted to determine how SPARC’s interaction with type IV collagen influences collagen within the BM. In *C. elegans* type IV collagen monomers are composed of two α1 chains encoded by *emb-9,* and one α2 chain encoded by *let-2* (71, 72). We examined if overexpression of SPARC alters the localization of a functional type IV collagen reporter, EMB-9::mCherry (henceforth referred to as collagen::mCherry (73)). We observed that overexpressing SPARC decreased BM collagen::mCherry levels by approximately 50% (Fig 3A). In contrast, SPARC overexpression did not significantly alter the levels of the other key BM scaffolding component laminin, suggesting overexpressed SPARC specifically regulates type IV collagen levels in the BM (Fig 3A).

**Figure 3.**
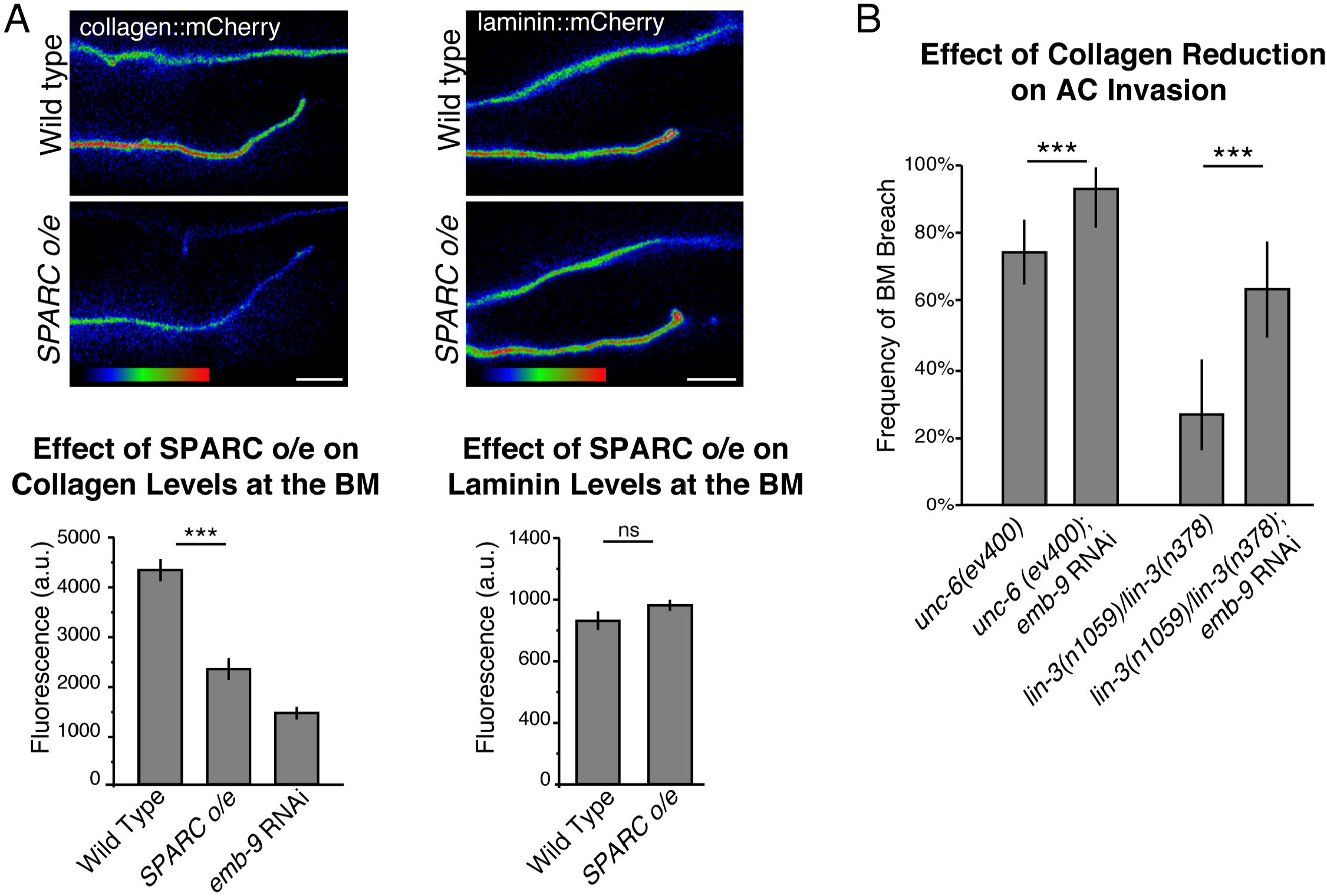
SPARC overexpression promotes AC invasion by decreasing type IV collagen levels in the BM. (A) A spectral representation of the fluorescence intensity shows collagen::mCherry (left) and laminin::mCherry (right) fluorescence at the BM in wild type (top) and *SPARC overexpression* (o/e; *syIs115;* bottom) animals. The graph below depicts the average levels of fluorescence for collagen::mCherry and laminin::mCherry at the BM (n≥16 for each treatment). (B) The graph depicts the frequency of BM breach in animals treated with RNAi targeting the *emb-9* collagen α1 subunit (n≥50 for each treatment). Error bars denote the standard error of the mean (SEM, A) or 95% confidence intervals with a continuity correction (B). ⋆⋆⋆ denotes p<0.0005 by unpaired two-tailed Student’s t-test (A) or two-tailed Fisher’s exact test (B). Scale bars denote 5 pm.

### Reduction of type IV collagen can account for SPARC’s pro-invasive activity

Given the key structural role of type IV collagen in BM, our observations suggested that overexpressed SPARC might promote invasion by reducing the levels of type IV collagen. To determine if reduction of collagen levels in the BM alone is sufficient to enhance AC invasion, we used RNAi targeting *emb-9* to decrease type IV collagen levels in the BM (an average of 63% reduction in collagen::mCherry fluorescence, mimicking overexpression of SPARC, Fig 3A). Similar to SPARC overexpression, decreasing the levels of type IV collagen significantly restored AC invasion in multiple mutant backgrounds where invasion is delayed or inhibited (Fig 3B). Together, these results suggest that SPARC promotes AC invasion by decreasing the level of type IV collagen in the BM.

### Overexpressed extracellular SPARC reduces collagen incorporation into the BM

SPARC and type IV collagen are both predominantly expressed in the *C. elegans* body wall muscle before being secreted and assembled in BMs throughout the animal (16, 67). Consistent with observations in *Drosophila* (40), SPARC and collagen colocalize in intracellular vesicles in *C. elegans* (S3 Fig). Much of the intracellular colocalization is likely rough ER, as SPARC::GFP colocalized with the ER marker TRAM::mCherry (S3 Fig (74)). Although colocalized during vesicle trafficking, previous studies have largely indicated that SPARC does not regulate intracellular trafficking of type IV collagen (12, 41). We thus sought to determine if overexpression of extracellular SPARC was sufficient to decrease BM type IV collagen. To test this, we separated the site of SPARC overexpression from the site of type IV collagen production. We took advantage of the neurons, which constitute a large population of cells in *C. elegans.* Neurons in *C. elegans* have not been reported to express SPARC but do express other ECM proteins, indicating that they are capable of secreting matrix proteins into the extracellular space (75). We drove SPARC using the pan-neuronal *rab-3* promoter ((76); *rab-3>SPARC::GFP).* Strikingly, we found that neuronally expressed SPARC decreased the amount of type IV collagen in the BM and promoted AC invasion (Fig 4). Notably, these strong effects were observed even though neuron-generated overexpression of SPARC::GFP resulted in less additional BM-localized SPARC than overexpressed SPARC::GFP driven by the endogenous SPARC promoter (S2 Fig). We conclude that overexpressed SPARC has an extracellular role in reducing type IV collagen levels in the BM.

**Figure 4.**
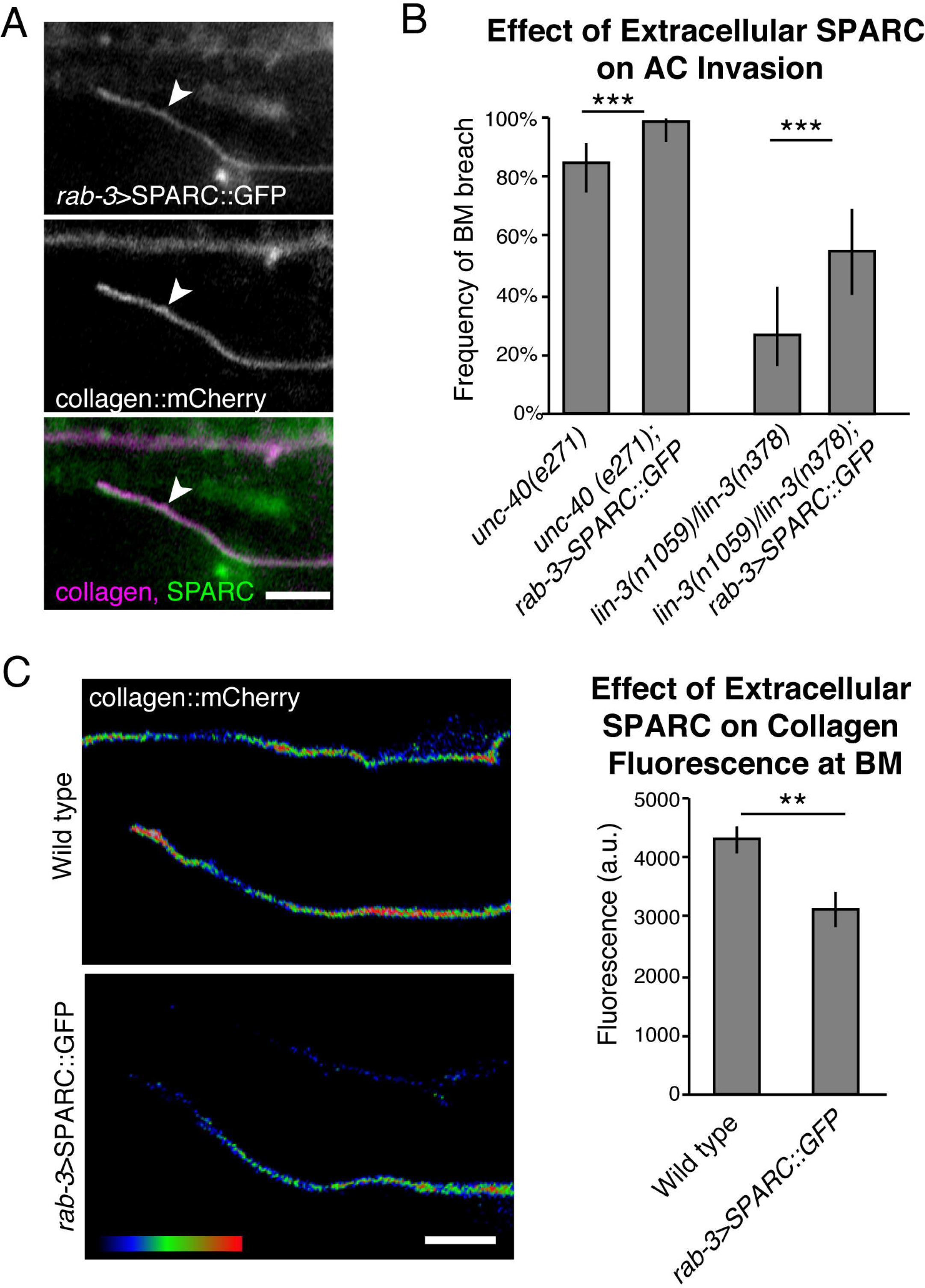
SPARC functions extracellularly to reduce BM type IV collagen levels. Neuronally expressed SPARC *(rab-3>SPARC::GFP;* top) is found at the BM (collagen::mCherry, center; merge, bottom). (B) Graph depicts the frequency of ACs breaching the BM in *unc-40(e271)* and vulvaless *(lin-3(n1059)/lin-3(n378))* animals when SPARC is overexpressed in the neurons *(rab-3>SPARC::GFP)* compared to worms not overexpressing SPARC (n≥49 for each genotype). ⋆⋆⋆ denotes p<0.0005 by two-tailed Fisher’s exact test, error bars show 95% confidence intervals with a continuity correction. (C) A spectral representation of collagen::mCherry fluorescence at the BM when SPARC is expressed neuronally is shown on the left and the quantification of collagen fluorescence at the BM is shown on the right (n=15 animals for each treatment). ⋆⋆ denotes p<0.005 by two-tailed unpaired Student’s t-test; error bars show SEM. Scale bars denote 5 pm.

### SPARC overexpression slows collagen incorporation into the BM

The lower levels of BM collagen in animals overexpressing SPARC suggested that extracellular SPARC might regulate the rate of collagen turnover (addition and/or removal) in the BM. To determine if SPARC overexpression alters collagen turnover, we examined fluorescence recovery after photobleaching (FRAP) of collagen::mCherry in the BM. We photobleached collagen::mCherry in the BM surrounding half of the uterine tissue and measured FRAP, using the unbleached half of the uterine tissue as a control to estimate total BM collagen levels throughout the experiment. Bleaching this large region rendered the extremely limited diffusion of BM localized type IV collagen negligible (Ihara et al., 2011). In wild type worms, 44±4% of the total collagen in the BM was replaced with fluorescent collagen two hours after photobleaching. In contrast, when SPARC was overexpressed under its endogenous promoter, only 25±3% of collagen recovered two hours after photobleaching (p<0.005 compared to wild type; Fig 5). These observations suggest that excessive extracellular SPARC decreases collagen addition to the BM. In sum, these results indicate that SPARC overexpression decreases type IV collagen incorporation into the BM, which weakens this barrier to cell invasion.

**Figure 5.**
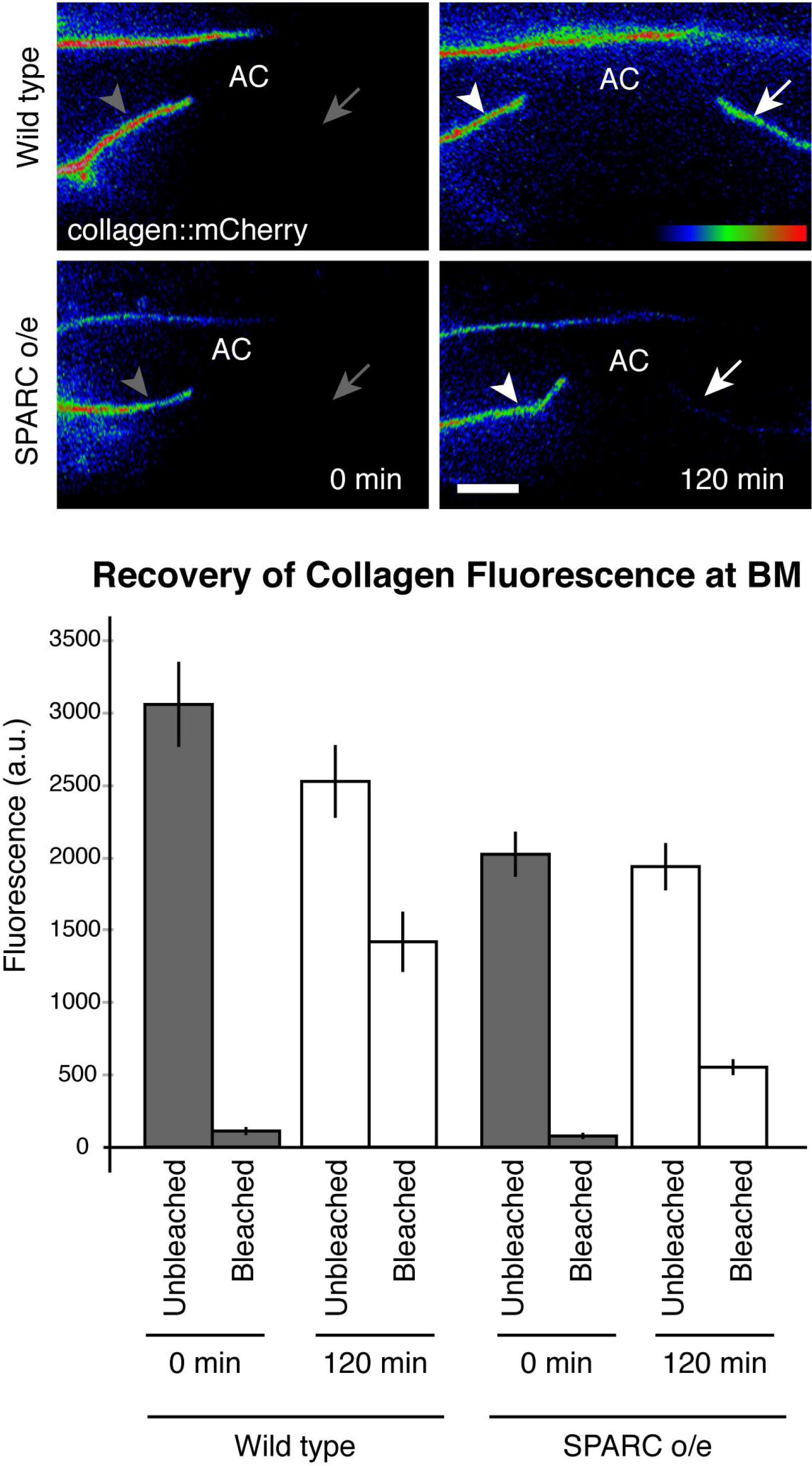
SPARC overexpression slows type IV collagen recovery in the BM. Half of the uterine tissue was photobleached by exposing collagen::mCherry within this region to 561 nm light for two minutes (0 min recovery, left; arrow denotes region of bleach, arrowhead denotes unbleached control region). Collagen::mCherry fluorescence recovery was assessed two hours later (120 min, right). Average collagen::mCherry fluorescence levels for the control and bleached regions at 0 min and 120 min is graphed below. An average of 44±4% of the total collagen recovered in wild type and 25±3% of collagen in worms overexpressing SPARC (n≥10 for each condition, p<0.005 by two-tailed unpaired Student’s t-test).

### Reduction of SPARC perturbs collagen trafficking during development

Our studies above suggested that increased SPARC expression interferes extracellularly with type IV collagen incorporation into the BM. Recent work in *Drosophila* has suggested that SPARC normally functions extracellularly to promote type IV collagen solubility after secretion (41). To investigate the normal role of SPARC in regulating type IV collagen assembly in *C. elegans,* we examined the type IV collagen-rich BM surrounding the pharynx, the muscular foregut of the worm that grinds food and transports it to the intestines (77, 78). The pharynx does not express type IV collagen, suggesting that collagen is recruited to the pharyngeal BM from a distant collagen production site, likely the body wall muscle (67). We reasoned that if SPARC acts to promote extracellular collagen solubility, then the pharyngeal BM would be particularly sensitive to reduction of SPARC function as it is localized at a distance from the site of type IV collagen synthesis (unlike the gonadal BM which can receive collagen from the nearby distal tip cell and sex muscles (67)).

Consistent with previous reports, we found that worms harboring putative null alleles in the SPARC locus *ost-1*(*tm966*) and *ost-1*(*tm6331*)) are embryonic and early larval lethal precluding analysis of SPARC function during later larval development (S4 Fig) (66). Thus, we reduced late larval SPARC levels by RNAi-mediated targeting (an average of 97% reduction, see Methods). SPARC reduction led to a dramatic decrease in type IV collagen levels in the pharyngeal BM (approximately 60% reduction in collagen::mCherry fluorescence compared to wild type animals, Fig 6A). This loss of collagen was frequently accompanied by severe morphological defects in the pharyngeal bulbs (n=20/33 animals, Fig 6A), indicating that the structural integrity of the pharynx had been compromised. Levels of the BM component laminin were not significantly affected by SPARC knockdown (Fig 6A), suggesting that SPARC is specifically required for type IV collagen incorporation into the BM. Notably, we also observed an approximately two-fold increase in accumulation of collagen at the surface of the body wall muscle (where type IV collagen is expressed and secreted) after reduction of SPARC (Fig 6B). Together, these results suggest that SPARC is required to transport type IV collagen from the surface of the body wall muscle to the pharyngeal BM during development.

Interestingly, we found that reduction of SPARC did not alter the levels of collagen surrounding the gonadal BM at the time of AC invasion (Fig 6C). As the distal tip cell of the somatic gonad and vulval sex muscles both produce type IV collagen and are in direct contact with the gonadal BM, SPARC may not be required to traffic extracellular collagen to the gonadal BM or very low levels of SPARC might be sufficient for transport. Consistent with unaltered collagen levels in the BM, SPARC reduction did not change the frequency of AC invasion in the *unc-40* (netrin receptor DCC) mutant background (Fig 6D). In addition, reduction of SPARC did not affect the timing or frequency of invasion in wild type animals (n=49/50 animals invaded by the P6.p four-cell stage and 50/50 by the P6.p eight-cell stage). Together, these results indicate that SPARC has an important role in regulating the transport of type IV collagen to tissues located at a distance from sources of collagen production. Furthermore, our results suggest that overexpression of SPARC disrupts its normal extracellular function in collagen trafficking, leading to a reduction in collagen BM deposition.

**Figure 6.**
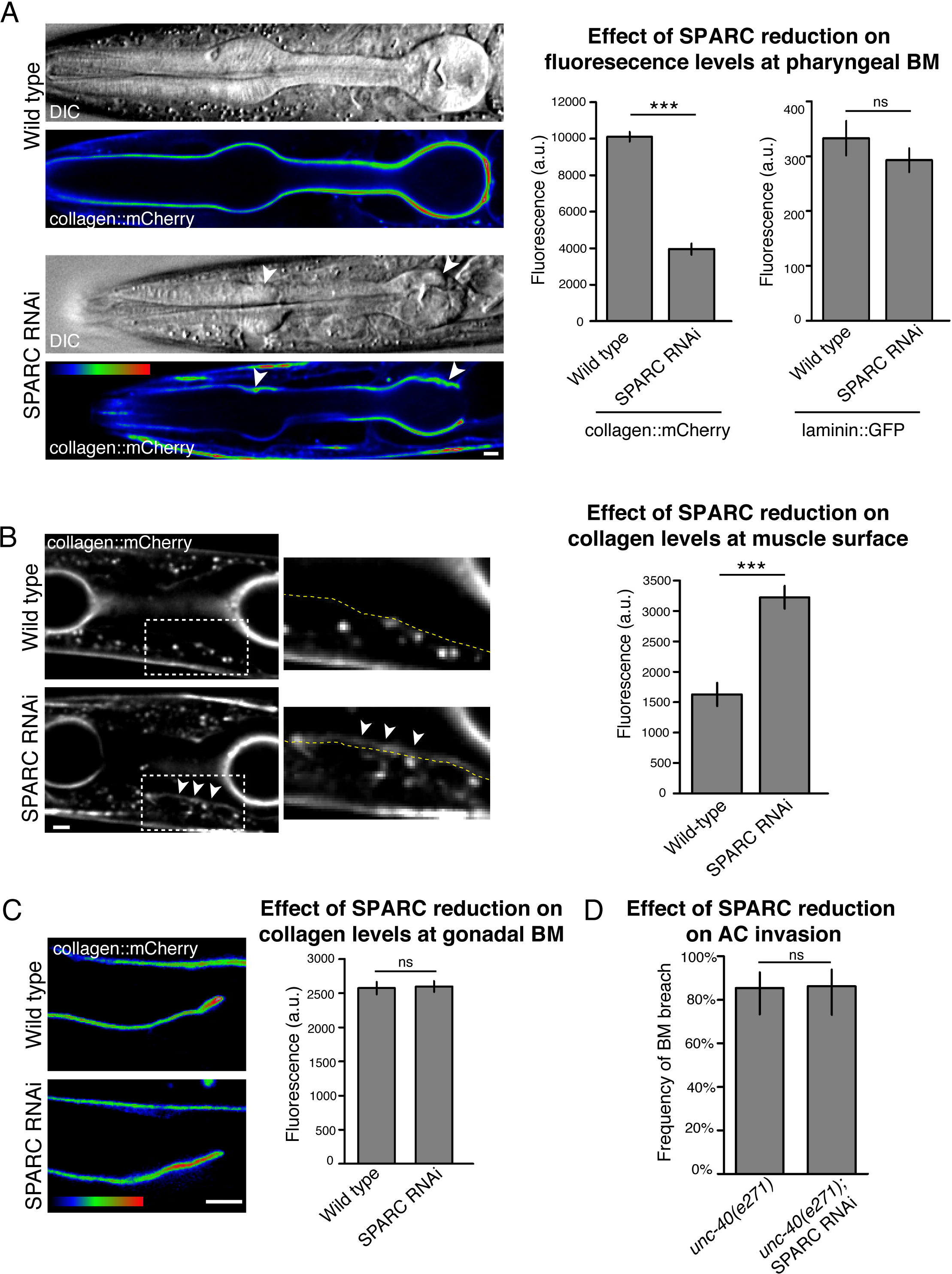
SPARC regulates the transport of type IV collagen during development. (A) DIC images and corresponding spectral representations of fluorescence intensity showing collagen::mCherry in the pharyngeal BM of a wild type animal (top panel) and SPARC RNAi-treated animal (bottom). Arrowheads indicate BM deformations after SPARC reduction. The average collagen::mCherry and laminin::GFP fluorescence levels at the pharyngeal BM in wild type and SPARC RNAi-treated animals are quantified in the graph on the right and compared using a two-tailed unpaired Student’s t-test (n>15 for each treatment; ⋆⋆⋆ denotes p<0.0005 for collagen::mCherry; p=0.3 for laminin::GFP; error bars show SEM). (B) Representative fluorescence images of collagen::mCherry in the Z-plane of body wall muscle cells in a wild type animal (top) and SPARC RNAi-treated animal (bottom). The boxed regions indicate body wall muscle cells and are magnified on the right with dashed lines representing the muscle surface as determined by corresponding DIC images. Collagen::mCherry accumulated abnormally at the muscle surface in the SPARC RNAi-treated animal (arrowheads), and was difficult to detect at the muscle surface in the wild type animal. The average collagen::mCherry fluorescence levels at the pharyngeal BM are quantified in the graph on the right (n=6 wild type and 12 RNAi treated animals). ⋆⋆⋆ denotes p<0.0005 by two-tailed unpaired Student’s t-test; error bars show SEM. (C) Spectral representation of collagen::mCherry fluorescence intensity at the gonadal BM in wild type (top) and SPARC RNAi treated (bottom) representative animals. The average collagen::mCherry fluorescence levels are quantified in the graph on the right (n>40 for each treatment; p=0.8 by unpaired two-tailed Students t-test; error bars show SEM). (D) Graph depicts the frequency of ACs breaching the BM in *unc-40(e271)* animals treated with SPARC RNAi (n>51 for each treatment; p=1.0 by two-tailed Fisher’s exact test). Error bars show 95% confidence intervals with a continuity correction. Scale bars denote 5 pm.

## Discussion

The matricellular protein SPARC has been implicated in regulating many molecular functions important in tissue growth, morphogenesis, and homeostasis; including cell-matrix adhesion, growth factor activity, and ECM dynamics (18, 79). The diverse functions of SPARC, compounded by the challenge of studying cell invasion and BM dynamics *in vivo,* have made it difficult to understand how overexpression of SPARC in metastatic cancers promotes cell invasion and tumor spread (15, 18). Using AC invasion in *C. elegans* as an *in vivo* model to examine the role of SPARC in regulating BM transmigration, our genetic, site of action and live imaging studies indicate that SPARC overexpression functions permissively to facilitate cell invasion by inhibiting type IV collagen deposition in the BM, thus weakening BM barrier properties (summarized in Fig 7).

**Figure 7.**
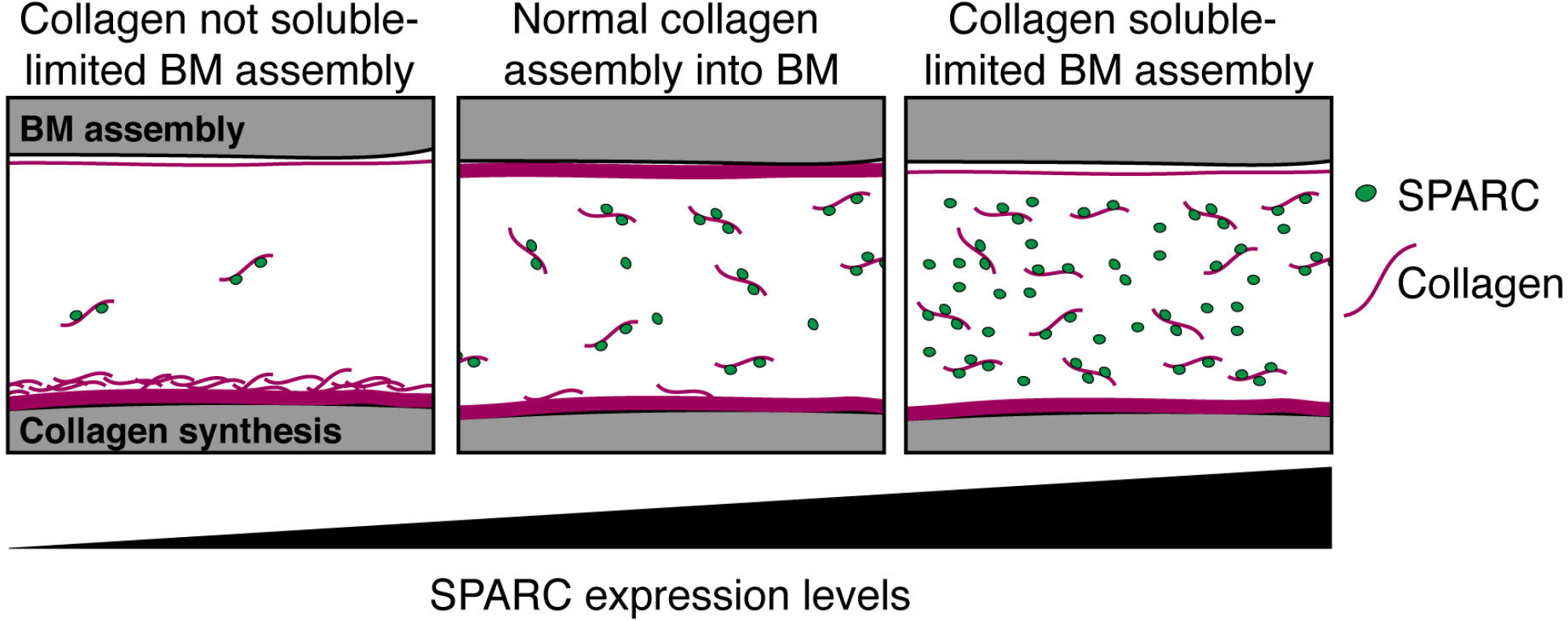
Model of SPARC function. Extracellular SPARC acts as a type IV collagen chaperone that maintains collagen solubility by inhibiting collagen polymerization and/or binding of collagen to cell surface receptors. When SPARC levels are low, type IV collagen solubility in the extracellular space is reduced resulting in an accumulation of collagen at the surface of collagen expressing cells and a decrease in type IV collagen reaching distant sites of BM assembly. At endogenous levels, SPARC helps transports collagen after secretion from sites of synthesis to distant BMs. When SPARC is overexpressed, collagen solubility is increased, resulting in reduced BM collagen incorporation. Previous studies have indicated that SPARC can bind collagen in multiple locations, with two or three SPARC molecules per collagen being the most common stoichiometry (44, 45).

While overexpression of SPARC did not alter the timing of the normal AC invasion program, we found that it dramatically restored AC invasion in numerous genetic backgrounds where AC invasion is delayed or completely blocked. Furthermore, type IV collagen imaging and FRAP analysis showed that SPARC overexpression led to a significant decrease in collagen deposition in the BM, and that RNAi-mediated reduction in BM collagen mimicked the proinvasive activity of SPARC. These data provide compelling evidence that SPARC overexpression promotes invasion by reducing BM type IV collagen levels. Changes in the structure and composition of ECMs can signal to invasive cells through integrin clustering or biomechanical pathways (4, 5); thus altering the properties of the BM may lead to an increase in the pro-invasive signals the AC receives from its environment. Alternatively, as the type IV collagen lattice is highly cross-linked and endows BM with its mechanical strength and barrier properties (4, 5), decreasing the levels of type IV collagen may weaken the BM, allowing this barrier to be more easily breached by invading cells. As we found that SPARC overexpression broadly promotes invasion after the reduction or absence of all previously identified signaling pathways regulating AC invasion (netrin, integrin, Fos and vulval cue pathways), our data support the idea that SPARC overexpression does not augment one of these previously identified pathways. Instead, we hypothesize that by limiting type IV collagen addition to BM, SPARC weakens the BM and allows it to be transmigrated more easily by ACs that have limited invasive capacity. Consistent with this idea, decreasing type IV collagen levels in BM has also been proposed to facilitate neutrophil extravasation through the venule BM (3, 80, 81). Furthermore, studies in colon cancer have suggested that reduction in type IV collagen expression via hypermethylation of collagen encoding genes may create a weaker BM that is more permissive to invasion (82).

The co-emergence of SPARC with collagen in basal metazoans and the presence of the highly conserved collagen-binding site in SPARC proteins (37, 83) suggest that SPARC and collagen have an ancient functional relationship. Previous work on *Drosophila* larvae has shown that in the absence of SPARC, collagen accumulates around the fat body (the site of collagen synthesis) and is absent from the BM of the ventral nerve cord, which does not express type IV collagen (12, 40, 41). We observed a similar accumulation of collagen at the surface of collagen-expressing body wall muscle cells upon reduction of SPARC in *C. elegans,* and a reduction of collagen within the pharyngeal BM, which does not express type IV collagen. These observations are consistent with the idea that SPARC acts to promote type IV collagen solubility, allowing it to be distributed from sites of secretion to the BM of distant tissues lacking its expression (Fig 7). Binding of SPARC to collagen might promote collagen solubility by preventing collagen from interacting with cell surface receptors and existing BM, or by inhibiting collagen polymerization (39, 41). In organs such as the somatic gonad that neighbor sites of type IV collagen synthesis, SPARC might not be required to move extracellular collagen or may play a more limited function in collagen trafficking. We suspect that when SPARC levels are high, collagen remains soluble and fails to incorporate as efficiently into BMs. The excess soluble collagen in the extracellular fluid is likely removed by coelomocytes, macrophage-like scavenger cells that endocytose numerous molecules in the body cavity fluid (84). SPARC may play a similar role in promoting collagen I solubility as cell culture studies have shown that addition of exogenous SPARC results in collagen I removal from a pre-assembled ECM (46). Together, our studies strongly support the emerging notion that SPARC acts as an extracellular type IV collagen chaperone (39, 41) whose levels play a crucial role in mediating collagen transport and deposition in ECMs.

Although previous studies have suggested that SPARC may either positively or negatively regulate laminin (40, 41, 85), we find that SPARC overexpression did not affect laminin deposition. This is consistent with previous findings during *Drosophila* development also showing that SPARC affects collagen but not laminin deposition (12, 86). As the structure and composition of BMs vary between tissues, the differing affect of SPARC on laminin may reflect variances in the structure and composition of BMs, or may be an indirect affect of altering collagen. Previous studies have also shown that SPARC interacts with collagen intracellularly to inhibit collagen deposition in the BM of *Drosophila* egg chambers, thus SPARC overexpression may alter collagen secretion (86). Our studies also showed that SPARC and collagen colocalized in the bodywall muscle, potentially in the ER, however, we found that extracellular SPARC was sufficient to alter collagen levels at the BM and promote invasion. While we cannot rule out the possibility that SPARC overexpression alters collagen secretion, the ability of extracellular SPARC *(rab-3>SPARC)* to recapitulate SPARC’s regulation of type IV collagen and invasion offers clear evidence that SPARC functions in the extracellular milieu to regulate collagen deposition in BM.

Suggesting an important and context-dependent relationship with cancer, both increased and decreased SPARC levels are associated with tumorigenesis (18, 87). A decrease in SPARC expression has been observed in numerous cancers including acute myeloid leukemia, multiple forms of lung cancer, pancreatic ductal adenocarcinoma, and ovarian carcinoma (88-92).

Adding exogenous SPARC to cell lines derived from these cancers inhibits cancer cell growth, suggesting that a loss of SPARC promotes tumorigenesis by facilitating cell proliferation (90, 92-94). Overexpression of SPARC is also associated with many cancers and high expression in the cancer cells often increases metastatic potential (26, 31, 64). Interestingly, in a subset of epithelial cancers increased SPARC expression is confined to epithelial cells surrounding the malignant cells (88, 89, 92, 95-99). Our observations indicating that extracellular SPARC reduces type IV collagen deposition suggests that increased SPARC expression from any nearby tissue would likely reduce BM type IV collagen levels of the growing tumor and tumor vasculature, making these tissues more readily traversed by malignant cells. We observed that a relatively small increase in SPARC expression enhances AC invasion, suggesting that SPARC expression may not need to be dramatically altered to promote metastasis. Given that our data suggest that excessive SPARC acts to weaken the BM barrier, SPARC overexpression alone may not be sufficient to endow a cell with metastatic capabilities, but may instead facilitate BM breach for a cell that already has a basal level of invasive ability. Supporting this idea, breast cancer lung metastases overexpressing SPARC require SPARC to promote invasion, but overexpressing SPARC is not sufficient to drive metastasis (21). Together our data suggests that SPARC expression must be tightly regulated to maintain ECM in a growing tissue, and that therapeutic strategies buffering SPARC activity to within the optimal range for BM collagen assembly may limit cell invasion during cancer metastasis.

## Materials and Methods

### *C. elegans* strain and culture information

*C. elegans* strains were cultured as previously described (100). The wild type strain was N2. In the text and figures, we refer to linked DNA sequences that code for a single fusion protein using a (::) annotation. For designating linkage to a promoter we use a (>) symbol. Alleles and transgenes used in this study are as follows: *syIs115(SPARC::GFP, unc-119(+)), syIsJJ3(SPARC::GFP, unc-119(+)), qyIs46(emb-9::mCherry, unc-119(+)), qyIs202(hsp>SPARC, myo-2>GFP, unc-119(+)), qyIs240(hsp> SPARC^R152L^'^Q159A^::GFP, unc-119(+), myo-2>GFP), qyEx480(Tram::mCherry), qyEx478(myo-3>mCherry::arf-6, myo-2>mCherry), qyIs432(rab-3>SPARC::GFP, unc-119(+)), qyIs128(lam-1::mCherry, unc-119(+));* LGI: *unc-40(e271);* LGII: *rrf-3(pk1426);* LGIII: *unc-119(ed4), ina-1(gm39);* LGIV: *lin-3(n1059)/lin-3(n378), ost-1(tm966), ost-1(tm6331), qyIs10(lam-1::GFP, unc-119(+));* LGV: *fos-1(ar105);* LGX: *unc-6(ev400)*.

### Microscopy, image acquisition, processing, and analysis

Images were acquired using a Hammamatsu EM-CCD camera, a spinning-disk confocal microscope (CSU-10; Yokogawa Corporation of America) mounted on an AxioImager base (Carl Zeiss) with 40× and 100× Plan Apochromat objectives (1.4 NA) and controlled by μManager (101). Worms were photobleached on a Zeiss AxioImager A1 microscope with a ×100 plan-apochromat objective by exposing a portion of the worms to 561 nm light for 2 minutes. Acquired images were processed to enhance brightness/contrast using ImageJ 1.40g and Photoshop (CS6 Extended Adobe Systems, InC., San Jose, CA). Colocalization analysis was performed on confocal z-using the “Coloc” module in IMARIS 7.4 (Bitplane, Inc., Saint Paul, MN). Measurements of collagen::mCherry intensity at the BM were collected in ImageJ by measuring the mean grey value of a 3 pixel width line drawn along the BM. An equivalent line was drawn in an adjacent region of the worm to measure the background signal within the worm, which was subtracted from the signal at the BM.

### Analysis of AC invasion

AC invasion was scored at the P6.p four-cell and eight-cell stage as previously described (53). Worms were staged by the divisions of the underlying P6.p vulval precursor cells. Animals were scored as “complete invasion” if the BM was removed beneath the AC, “partial invasion” if the BM was breached but not cleared beneath the AC, and “no invasion” if there was no BM breach.

### Generation of SPARC overexpression lines

SPARC::GFP strains were made by injecting pOST8-X1/2 at 50-100 ng/μl with *unc-119(+)* into *unc-119(ed4)* and then integrated. *qyIs202* was made from a plasmid containing *hsp>SPARC::GFP,* a gift from J. Schwarzbauer, created by fusing a 349 bp segment containing the 1.48 intergenic region of hsp16-1/48 to the full *ost-1* open reading and an additional 800 bp 3’ flanking region. *hsp> SPARC^R152L^’^Q159A^* was made by inserting GFP between the first and second exons of *hsp>SPARC* followed by site directed mutagenesis. *rab-3>SPARC::GFP* was assembled by inserting the following regions into pBlueScript: *rab-3* promoter (Kpn1, Apa1), ost-1 exon1::GFP from pOST8-X1/2 (Apa1, EcoRV) and ost-1 exon 2-6 from cDNA (EcoRV, Not1).

### Quantitative Real-Time PCR

Total RNA was prepared from mixed-staged populations (TriReagent, Sigma) and cDNA synthesized using random primers (Invitrogen). An ost-1-specific primer set was designed as follows: forward primer 5’GCATGGCTGACTGGCTCTT3’, reverse primer 5’GTGGAGCTCACGTCTCTTCTT3’, and FAM-labeled reporter with non-fluorescent quencher spanning exon 4/exon 5 boundary 5’CCAGGTCATGAAGGAGCTT3’. *act-4* (actin) was selected as the normalization gene with FAM-labeled reporter with non-fluorescent quencher 5’GGAGACGAAGCCCAGTCCAAGAGAG3’ (Assay ID Ce02508047_s1, Applied Biosystems, Foster City, CA). QPCR reactions used 1ng cDNA, Taqman Chemistry, and an ABI prism 7000 (Applied Biosystems). Relative quantification of *ost-1* mRNA levels were standardized against *act-4* levels and normalized to wild type N2. Data analysis was performed using methods as described in the ABI Prism 7700 user bulletin #2. Each biological sample was performed in duplicate and the PCR reaction was repeated in triplicate.

### RNA interference

*ost-1* cDNA fragments 67-660 bp and 129-727 bp were amplified by PCR from N2 cDNA, cloned into the L4440 (pPD129.36) vector, and transformed into HT115 bacteria. Synchronized, L1-arrested larvae were grown on regular OP50 bacteria at 20°C to the L4, after which they were transferred to *ost-1* RNAi plates. By performing an L4 RNAi plating, we were able to achieve ˜97% reduction of SPARC as measured by SPARC::GFP levels in L4 progeny (L4440 treated animals fluorescence at BM= 6900±700 a.u.; *ost-1* RNAi treated animals fluorescence at BM=200±100 a.u.; ±SEM; n≥20 animals). SPARC reduction phenotypes were assessed in L3/L4 progeny (at the time of AC invasion, for gonadal BM measurements) or young adult progeny (for pharyngeal BM measurements). RNAi targeting *emb-9* was acquired from the Ahringer library (102), fed to synchronized L1 larvae, and phenotypes were assessed at the L3/L4 at the time of invasion. The empty vector L4440 was used as a negative control in all RNAi experiments. All RNAi vectors were sequenced to verify the correct insert.

### Immunocytochemistry and western blot analysis

Worm strains were sectioned and immunostained as described and incubated with anti-LET-2 (NW68, 1:200)(67), anti-SPARC (1:400)(69) or anti-EPI-1 (1:500)(103). Worms were prepared for western blot by washing synchronized L4 populations off NGM agar plates with OP50. Worms were washed with M9 3-5 times and allowed to digest the bacteria remaining in their gut for 30 minutes before flash freezing in liquid nitrogen. Worms were boiled in GLB buffer (2% SDS, 10% glycerol, 50mM Tris-Ph 6.8, 5% β-mercaptoethanol) for 10 minutes, sonicated for 10 seconds, and boiled a final time immediately before loading onto a 4-15% MiniPROTEAN TGX gel (BioRad, Hercules, CA). Blots were blocked with 5% dry milk in TBS-T and probed with anti-SPARC (1:100, (16) and E7 (1:250, Developmental Studies Hybridoma Bank) at 4°C overnight. Blots were analyzed using the ImageJ gel analyzer plug-in.

### Statistical Analysis

Statistical analysis was performed in JMP version 9.0 (SAS Institute) or Microsoft Excel, using either a two-tailed unpaired Student’s *t* test or a two-tailed Fisher’s exact test. Confidence intervals reported are 95% confidence intervals with a continuity correction. Figure legends specify which test was used.

## Acknowledgements

We thank J. Kramer and J. Schwarzbauer for strains and discussions, and M. Clay and M. Keenan for comments on the manuscript. The SPARC deletion alleles were provided by the *C. elegans* deletion mutant consortium (104). Some strains were provided by the Caenorhabditis Genetics Center, which is funded by NIH Office of Research Infrastructure Programs (P40 OD010440). 501

## Supporting Information Captions

**Figure S1. Fig Quantification of SPARC overexpression** (A) Quantitative Real Time PCR of SPARC levels in wild type animals and animals expressing integrated transgenes overexpressing SPARC::GFP *(syIs113* and *syIs115).* SPARC mRNA levels were standardized using *act-4* (actin) mRNA as a control and normalized to wild type (N2) levels. Error bars denote standard deviation in each of three replicates of the RT-PCR reaction. (B) A representative western blot shows endogenous SPARC (27 kD) and overexpressed SPARC::GFP (54 kD) in *sysIs115* and *syIs113* transgenic lines. An average of three western blots showed that *syIs115* animals express 2.5±0.4 fold more SPARC protein relative to endogenous SPARC and *syIs113* animals have a 5.4±1.6 fold increase in SPARC protein expression.

**Figure S2. Fig SPARC^R152L,Q159A^::GFP, rab-3>SPARC::GFP and SPARC::GFP localization at the BM.** (A) *hsp*>SPARC^R152L,Q159A^::GFP (green, left; overlaid with DIC, right) is found in the BM (arrows). (B) Quantification of SPARC::GFP fluorescence at the BM in SPARC overexpression lines. As SPARC overexpression does not appear to affect the expression of endogenous unlabeled SPARC (see S1 Fig), these graphs depict only the population of SPARC present in excess of endogenous levels. Error bars denote SEM. Scale bars denote 5 μm.

**Figure S3. Fig SPARC and type IV collagen colocalize in the ER.** (A) SPARC::GFP *(syIs115* line; left) and collagen::mCherry (center) colocalize in vesicles within the body wall muscle (overlay, right; average Pearson’s correlation coefficient=0.70 ± 0.01; n=8 animals). (B) Overexpressed SPARC::GFP *(syIs115;* left) is predominantly localized in the rough ER (as marked with TRAM::mCherry, middle; overlay, right; n=5 animals). Arrows highlight regions of colocalization.

Figure S4. Fig Characterization of two SPARC deletion alleles. Two non-overlapping deletions in the SPARC open reading frame *(ost-1)* were obtained from the *C. elegans* Gene Knockout Consortium. The location of the deletions *tm6331* and *tm966* is shown on top. Below, the progeny from a timed egg lay of *ost-1 (tm6331)/+* (left) or *ost-1 (tm966)/+* (right). The progeny of the egg lay were tracked for one week and their phenotypes were recorded. More than 25% of the progeny of *ost-1(tm6331)* and *ost-1(tm966)* did not survive to adulthood (green), suggesting that homozygotes of both mutant alleles are embryonic (dark red) or larval (light red) lethal.

